# The contribution of striatal pseudo-reward prediction errors to value-based decision-making

**DOI:** 10.1101/097873

**Authors:** Ernest Mas-Herrero, Guillaume Sescousse, Roshan Cools, Josep Marco-Pallarés

**Author notes:** Both authors equally contributed to this work (Ernest Mas-Herrero, Guillaume Sescousse). **Corresponding Author:** Ernest Mas-Herrero & Josep Marco-Pallarés, Montreal Neurological Institute, McGill University, Montreal, QC H3A 2B4, Canada.

## Abstract

Most studies that have investigated the brain mechanisms underlying learning have focused on the ability to learn simple stimulus-response associations. However, in everyday life, outcomes are often obtained through complex behavioral patterns involving a series of actions. In such scenarios, parallel learning systems are important to reduce the complexity of the learning problem, as proposed in the framework of hierarchical reinforcement learning (HRL). One of the key features of HRL is the computation of pseudo-reward prediction errors (PRPEs) which allow the reinforcement of actions that led to a sub-goal before the final goal itself is achieved. Here we wanted to test the hypothesis that, despite not carrying any rewarding value *per se*, pseudo-rewards might generate a bias in choice behavior when reward contingencies are not well-known or uncertain. Second, we also hypothesized that this bias might be related to the strength of PRPE striatal representations. In order to test these ideas, we developed a novel decision-making paradigm to assess reward prediction errors (RPEs) and PRPEs in two studies (fMRI study: n = 20; behavioural study: n = 19). Our results show that overall participants developed a preference for the most pseudo-rewarding option throughout the task, even though it did not lead to more monetary rewards. fMRI analyses revealed that this preference was predicted by individual differences in the relative striatal sensitivity to PRPEs vs RPEs. Together, our results indicate that pseudo-rewards generate learning signals in the striatum and subsequently bias choice behavior despite their lack of association with actual reward.

## Introduction

Reinforcement Learning (RL) theories have provided invaluable insights into reward-guided learning and decision-making. A core feature of RL theories is that actions are reinforced according to reward prediction errors (RPEs), that is, according to whether obtained outcomes are better or worse than expected (Sutton and Barto, 1998). It is now widely accepted that dopaminergic neurons projecting to the striatum play a pivotal role in the computation of RPEs (Schultz et al., 1997).

Although the computational principles of standard RL theories have accounted for a wide range of experimental findings in simple learning paradigms (Niv et al., 2002; Behrens et al., 2007), they often lead to sub-optimal behavior in complex situations involving a series of actions (Botvinick et al., 2009; Botvinick, 2012). For instance, when attending a conference in a new town, a series of actions is required to get from the hotel to the conference venue. If you get lost, you will need to think back and identify which of these actions need(s) to be adjusted next time. As the number of required actions increases, this operation becomes increasingly difficult for standard RL algorithms. One of the computational approaches that has emerged to solve this problem is reducing the number of actions through temporal abstraction as proposed by hierarchical reinforcement learning (HRL) algorithms (Botvinick et al., 2009; Behrens and Jocham, 2011; Frank and Badre, 2012; Badre and Frank, 2012; Botvinick et al.,2012).

In HRL, sequences of actions (e.g. exit hotel, turn right, go straight and turn left) are packaged together into subroutines evaluated according to their own subgoals (e.g. reach bus stop). Attaining a subgoal constitutes a so-called *pseudo-reward*, and any deviation from subgoal expectations generates a pseudo-reward prediction error (PRPE). Thus, in HRL there are two value functions – tracking rewards and pseudo-rewards – that are learnt in parallel via the same learning rule. HRL reduces the complexity of the learning problem by (1) focusing on a small set of decisions (the number of subroutines) rather than a large sequence of primitive actions and (2) confining behavioral adjustment to the actions within one subroutine when subgoal expectations are not met (Figure 1). In addition, PRPEs reinforce actions leading to the subgoal even before the final reward is obtained, therefore speeding up learning. fMRI studies have shown that both RPEs and PRPEs are computed in the ventral striatum (Ribas-Fernandes et al., 2011; Diuk et al., 2013).

**Figure 1:**
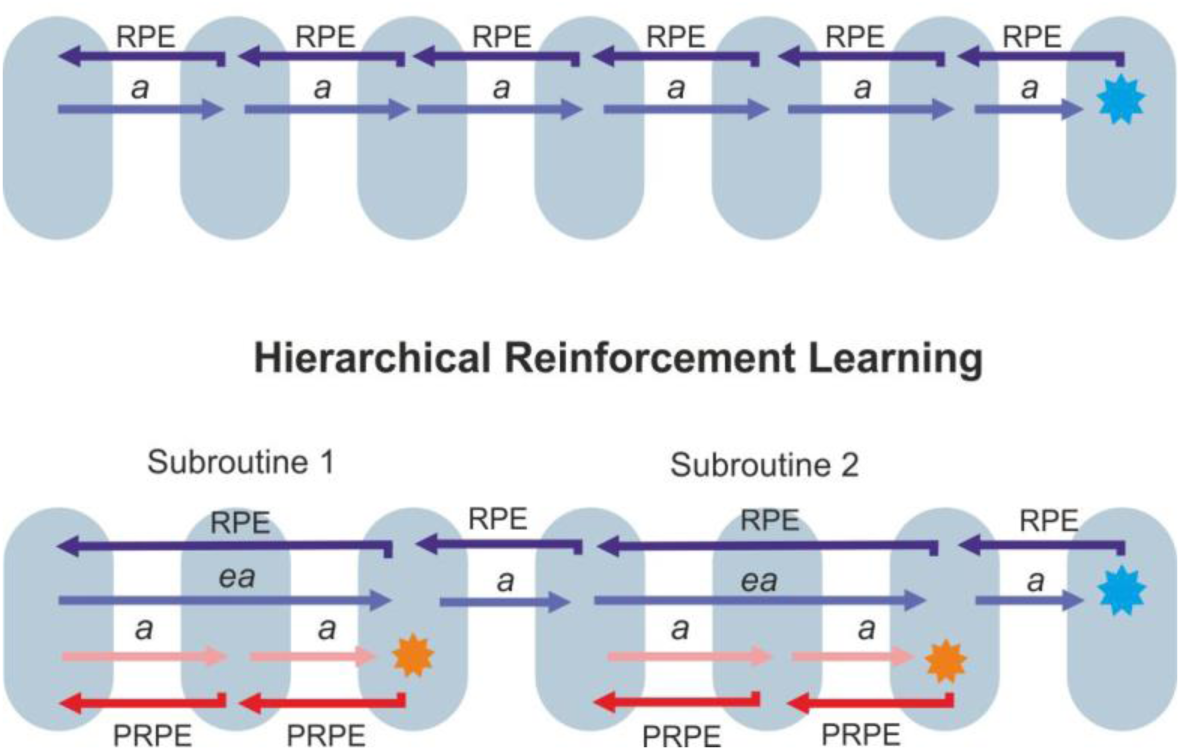
Main differences between classic Reinforcement Learning, as implemented in Temporal Difference (TD) learning models, and Hierarchical Reinforcement Learning (HRL). TD models aim to predict all events occurring in between an initial primitive action (*a*) and the final reward (blue star). All these events generate RPEs which are back-propagated in time. In order to accelerate learning, HRL algorithms divide primitive actions into subroutines of extended actions (*ea*) leading to subgoals/pseudo-rewards (orange star). RPEs are computed following the attainment of each subgoal/pseudo-reward. In doing this, RPE signals are back-propagated more rapidly (in the above example, 6 jumps are needed in TD learning to reach the initial action from the final outcome, while only 4 jumps are required in HRL). Importantly, HRL includes a second value function tracking the attainment of subgoals/pseudo-rewards. This value function uses PRPEs to reinforce those actions leading to a certain subgoal. The goal of the present study is to examine the parallel neural computation of RPEs and PRPEs and how they influence choice behavior.

However, the reinforcing properties of pseudo-rewards may also lead to ‘negative transfer’, i.e. preference for non-beneficial pseudo-rewarding options. Going back to the previous example, you might prefer to take a bus in front of your hotel followed by a 10 minute walk to the conference venue, rather than a bus that would first require a 10 minute walk but would then drop you off even more rapidly at the conference venue. This is because in the first option the first subgoal (taking the bus) is attained more rapidly and thus associated with more pseudo-reward. Such biases have been suggested on the basis of computational modeling (Botvinick et al., 2009), but have not been studied experimentally.

Here, we aimed to study how individuals develop such irrational biases towards pseudo-rewards and whether these depend on striatal sensitivity to RPEs and PRPEs. We developed a novel fMRI task in which participants had to collect as much money as possible from locked boxes. In order to collect the money, participants first had to unlock these boxes (subgoal). Once unlocked (pseudo-reward), each box could contain money (reward) or not. Crucially, participants had to choose between two keys to unlock the boxes. These two keys eventually led to the same amount of money but, by design, one of them unlocked more boxes than the other. We hypothesized that participants would be biased towards the latter, associated with more pseudo-reward, even though this decision was not related to higher monetary gains. In addition, if the striatum computes both RPEs and PRPEs, we hypothesized that the relative striatal sensitivity to RPEs vs PRPEs would predict participants’ bias.

## Materials and Methods

### Participants

Twenty-three students from the University of Barcelona (M = 21.9 years, SD = 2.7, 10 men) participated in the fMRI experiment. All participants were paid 20€ per hour and a monetary bonus depending on their performance. All participants gave written informed consent, and all procedures were approved by the local ethics committee.

### Experimental procedure

We developed a novel learning task inducing both reward and pseudo-reward prediction errors (Figure 2). Participants’ main goal was to accumulate as much money as possible. The money was placed into two boxes which were locked. Each box could only be unlocked with its own key. Thus, in each trial, the participants had to choose between two keys, in order to unlock the padlock and collect the money in the case the box contained some. Unlocking the box can be defined as a pseudo-reward as it is a non-rewarding state that constitutes a bottleneck to reach the reward. Crucially, one box could be unlocked 70% of the time but only contained money 30% of the time, while the other box could only be unlocked 30% of the time but contained money 70% of the time. Thus, the two boxes were associated with the same final reward probability (70% × 30% = 21%), but one of them was associated with a higher pseudo-reward rate (70% vs 30%). With this design we aimed to study whether pseudo-rewards may bias participants’ decision-making towards a preference for the most pseudo-rewarding option, despite the fact that this option did not confer any advantage in terms of monetary reward (Figure 2).

**Figure 2:**
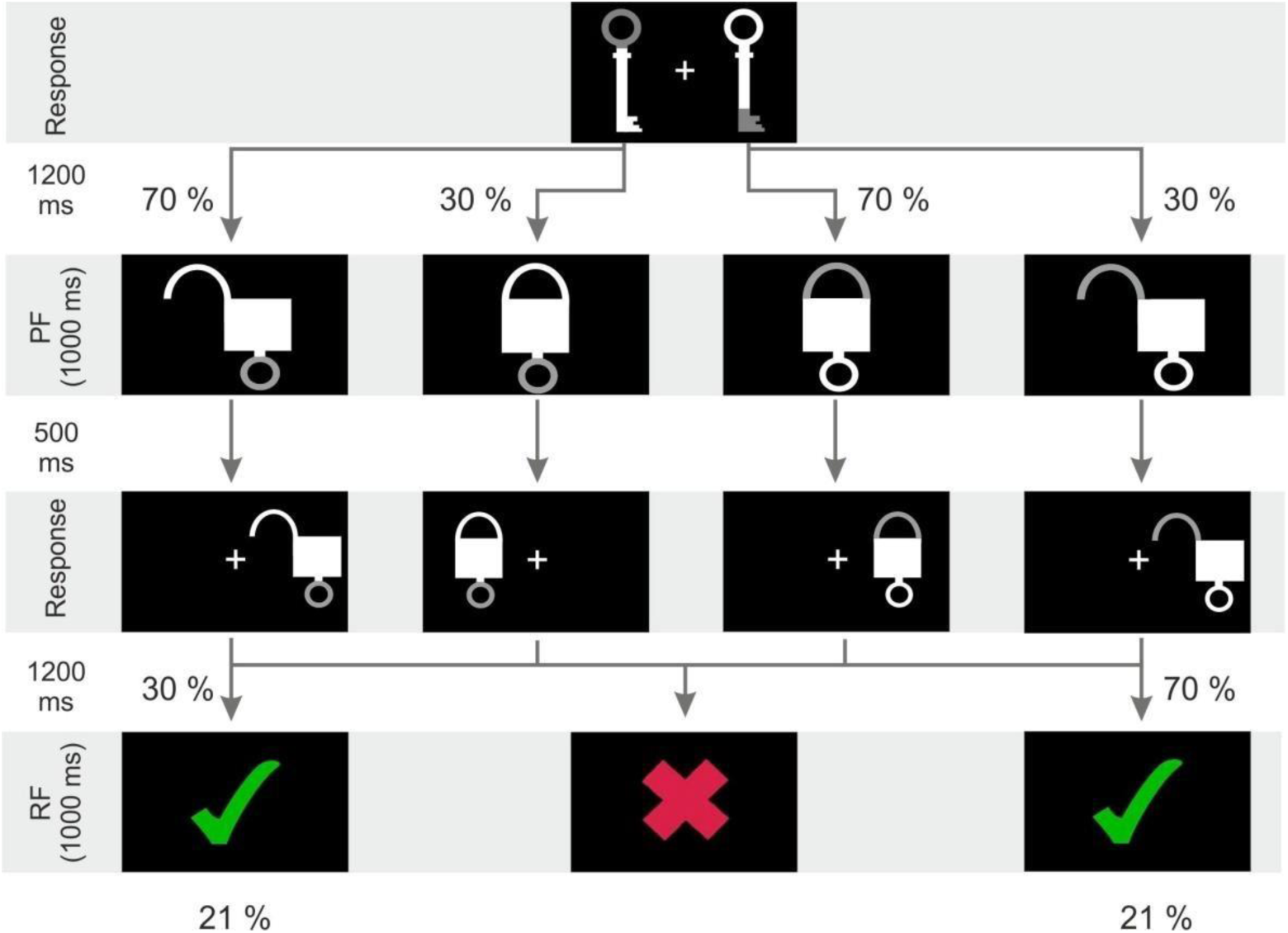
Task design. In the first-stage participants have to choose between two different keys. Each key unlocks a different type of box. In the second-stage a locked or unlocked padlock is presented depending on whether participants successfully unlocked the box or not. After using the key, and before the outcome is presented, participants have to remove the key from the padlock by indicating the position of the lock (third stage). Finally, either a green tick (reward) or a red cross (no-reward) is presented. The key on the left unlocks its associated boxes 70% of the time but these boxes contain money only 30% of the time. In contrast, the key on the right unlocks its associated boxes only 30% of the time but these boxes contain money 70% of the time. Overall, both keys are equally rewarded (21% of trials). *RF = Reward Feedback; PR = Pseudo-Reward feedback*.

Participants had to select, in less than 2 seconds, one of the two keys by pressing either the left (index finger) or right button (middle finger) of a response pad. If participants did not respond in time, a question mark appeared in the center of the screen for 1000 ms (these trials were discarded from the analyses). After a delay (1200 ms), a ‘pseudo-feedback’ indicated whether the padlock was unlocked or not (1000 ms). Then, 500 ms after the pseudo-feedback offset, the padlock was presented either on the left or the right side of the screen. Participants were told that they had to ‘remove the key’ from the padlock by indicating the position of the padlock (left or right button). Participants were asked to perform this action as fast as possible (within a time limit of 1000 ms), regardless of whether it had been unlocked or not. If they did not respond within the time limit, the final outcome was always negative. If they were successful, a ‘reward feedback’ was presented after a delay of 1200 ms, indicating whether the box contained money or not (green tick = 0.25€, red cross = 0€, 1000 ms). In trials in which the key was not properly removed or the padlock remained unlocked, a red cross indicating that the participant did not accumulate money was also presented in order to maintain the same structure in all trials (this ‘non-informative’ feedback was not further analyzed). After reward feedback a fixation cross remained on the screen until the end of the trial (mean trial duration was set to 11 seconds) followed by variable inter-trial interval (randomly jittered between 100 and 2500 ms, with 300 ms increments). Thus, participants were rewarded only if the following requirements were met: the box was unlocked, the key was properly removed in time and the box contained money. In other cases, the reward feedback was always a non-rewarding outcome. As mentioned previously, one of the keys (the most pseudo-rewarding key, referred to as PS+ key) had a greater probability of unlocking its associated boxes (p = 0.7) but those boxes had a low probability of containing money (p = 0.3). The other key (referred to as PS-key) presented the opposite pattern: it unlocked a smaller fraction of boxes (p = 0.3) but these boxes were more likely to contain money (p = 0.7). As a consequence both keys were equally rewarded (p = 0.21). Participants were told that a monetary bonus proportional to the amount of money accumulated during the task would be paid out at the end of experiment.

The task consisted of three blocks of 69 trials. Each block comprised of 46 forced-choice trials and 23 free-choice trials that were intermixed. Forced-choice trials were similar to the free-choice trials explained above with the difference that only one key appeared in the screen, either in the left or right side (random order). Participants had to press the corresponding button to select it. In half of the forced-choice trials, participants were forced to select the most pseudo-rewarding key. In the other half, they had to select the least pseudo-rewarding key. These forced-choice trials were introduced to ensure that participants would sample both options equally. In the remaining free-choice trials, participants could freely choose between the two different keys.

### Behavioral analysis

In order to study the impact of pseudo-rewards on participants’ choices we used two behavioral measures. First, we quantified the preference for the PS+ key, by computing the proportion of free-choices in which participants selected the PS+ key over the PS-key. To illustrate the effect of learning, we divided free-choice trials into 11 bins of 6 trials each, and plotted the preference for the PS+ key across bins. A repeated-measures ANOVA with time (i.e. bins) as a within-participant factor was performed to assess whether individuals’ preferences remained stable through the task or changed with learning.

Second, we performed a modified logistic regression to predict participants’ choices based on both the amount of reward and pseudo-reward obtained in previous trials (following a similar procedure as in Diuk et al., 2013). Specifically, we regressed participants’ choices onto the linear combination of rewards and pseudo-rewards obtained in the last four trials in which PS+/PS-choices were made. For each trial, we first computed the value of each key as follows:

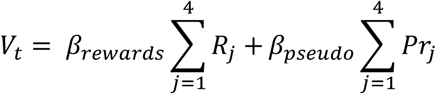

Where *R*_*j*_ was +1 or 0 depending on whether the participant received money or not on the *j*th-to-last time (s)he selected that key, and *Pr*_*j*_ was +1 or 0 depending on whether the participant was pseudo-rewarded (unlocked the box) or not the *j*th-to-last time (s)he selected that key. Participants’ free-choices were then logistically regressed on these values using a softmax equation:

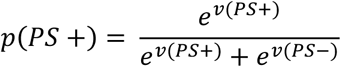

We optimized the two *β* parameters by minimizing the negative log likelihood using the *fminunc* function in MATLAB. Then, we tested whether the estimated parameters were significantly different from 0 at the group level using one-sample t-tests. The two *β* parameters represent the respective weights of past rewards and pseudo-rewards in participant’s choices. If participants are guiding their behavior based on the availability of both rewards and pseudo-rewards we would expect both *β*s to be positive and significantly different from 0.

### Reinforcement Learning Model

In order to estimate RPEs and PRPEs for the fMRI analysis, we implemented a temporal difference learning (TD) model in which two actions values are computed in parallel based on the history of rewards (Vr) and pseudo-rewards (Vpr), respectively (Diuk et al., 2013). Vr is calculated according to reward prediction error. In particular, two RPEs are computed in the current task: when the pseudo-feedback is presented (RPE1) and when the final outcome is reached (RPE2). In contrast, Vpr is calculated according to PRPEs. PRPEs only arise when the pseudo-feedback is presented and indicates whether the sub-goal has been accomplished or not. Thus, at the time of the pseudo-feedback two distinct prediction errors are computed, the RPE1 and the PRPE, related to the attainment of rewards and pseudo-rewards, respectively.

**RPE**. The action value *Vr* of each key (in the example below, the PS+ key) was updated when the pseudo-feedback was presented:

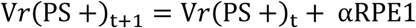

Where *α* is the learning rate, the subscript *t* represent number of trial, and RPE1 is the RPE term generated following the first action:

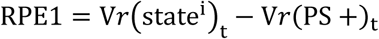

That is, RPE1 is generated by comparing the value of the state presented just after the selection of a key 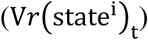 and the action value of that key (V*r*(PS +)_t_). In the current task four different states may be presented after the key selection: PS+ box unlocked, PS+ box locked, PS-box unlocked, PS-box locked (represented by *j* superscript). These states acquire value after the final feedback is presented trough a classic TD rule:

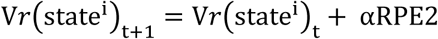

Where RPE2 is the prediction error term computed after the second action:

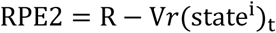

Where R was +1 or 0 depending whether the box contained money or not.

**PRPE**. PRPE were computed after each pseudo-feedback as:

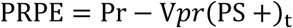

Where Pr was +1 or 0 depending on whether the box was unlocked or not on that trial, and *VPr*(PS+)_t-1_ is the expectancy of obtaining a pseudo-reward from the PS+ key. V*pr* represents action values according to the attainment of pseudo-rewards only. These action values are learned following the same rule than V*r*:

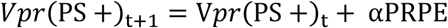

The learning rate (α) was set to 0.5 (Seymour et al., 2004; Wilson and Niv, 2015).

### fMRI data acquisition

fMRI data was collected using a 3T whole-body MRI scanner (General ElectricMR750 GEM E). Conventional high-resolution structural images (MP RAGE sequence, repetition time (TR) = 4.7 ms, echo time (TE) = 4.8ms, inversion time 450 ms, flip angle 12°, 1 mm isotropic voxels) were followed by functional images sensitive to blood oxygenation level-dependent (BOLD) contrast (echo planar T2*weighted gradient echo sequence, TR=2500 ms, TE=29 ms, flip angle 90°). Three functional runs were acquired, each consisting of 316 whole-brain volumes (40 slices, 3.1 mm in-plane resolution, 3.1 mm thickness, no gap, positioned to cover all but the most superior region of the brain and the cerebellum). In order to reduce susceptibility artifacts in the orbitofrontal cortex and the anterior parts of the ventral striatum, slices were orientated with an angle of 30 degrees with the plane intersecting the anterior and the posterior commissures (Weiskopf et al., 2006).

### fMRI data analysis

Pre-processing was carried out using Statistical Parametric Mapping software (SPM8, Wellcome Department of Imaging Neuroscience, University College, London, UK, www.fil.ion.ucl.ac.uk/spm/). Functional images were first sinc interpolated in time to correct for slice timing differences and were spatially realigned. Realigned images were then spatially smoothed with a 4 mm FWHM kernel before they were motion-adjusted using the ArtRepair toolbox (Mazaika et al., 2007). Specifically, functional images with significant motion artifacts were identified based on scan-to-scan motion (head position change exceeding 0.5 mm or global mean BOLD signal change exceeding 1.3%) and replaced by linear interpolation between the closest non-outlier volumes. Three participants were removed from the analysis as they presented more than 20% of volumes with artifacts in one or more runs due to excessive motion (see Mazaika et al., 2007) and, thus, the final sample consisted of 20 participants. In the remaining participants, 0.72% (SD = 1.11) of volumes were corrected for excessive motion on average. Then, the bias-corrected structural image was coregistered to the mean functional image and segmented by means of the Unified Segmentation implemented in SPM8. The resulting normalization parameters were applied to all functional images. Finally, functional images were spatially smoothed with a 7 mm FWHM kernel (the two-step smoothing of 4 mm and 7 mm is roughly equivalent to an overall smoothing of 8 mm, see Mazaika et al., 2007).

For the statistical analysis an event-related design matrix was specified. Seven regressors were included for each trial: key presentation, response, pseudo-feedback, presentation of the padlock to the left or the right, response and informative or non-informative feedback. Each response regressor was associated with a parametric regressor indicating whether the response was given with the middle or index finger. The pseudo-feedback regressor was associated with two parametric regressors modelling RPE1 and the PRPE computed at the time of the pseudo-feedback. The feedback regressor was associated with a parametric regressor modelling the reward prediction error computed at the time of the final outcome (RPE2). In order to achieve a uniform scaling of the output regression parameters, the parametric regressors representing RPE1, PRPE and RPE2 were standardized to a mean of 0 and a standard deviation of 1 (Erdeniz et al., 2013). The two parametric regressors included in the pseudo-feedback regressor were moderately correlated (M = .61). In order to avoid multicollinearity issues, the PRPE regressor was orthogonalized with respect to the RPE1. With this procedure we aimed to examine whether variations in PRPEs may still account for variations in the BOLD signal after removing the variance shared with RPE1. All regressors were subsequently convolved with the canonical hemodynamic response function and entered in a first level analysis. A high-pass filter with a cut-off of 128 s was applied to the time series. Three main contrasts of interest, testing the slopes of RPE1, PRPE and RPE2 regressors, were built at the first level. These contrast images were introduced into separate second-level group analysis based on one-sample t-tests. In order to study the conjunction of these three contrasts, first-level images were also introduced into a flexible factorial design. The conjunction analysis was formulated as a conjunction null hypothesis (Friston et al., 2005; Nichols et al., 2005) and should therefore only yield activations that were significantly present in all the contrasts introduced in the conjunction. Finally, in order to assess the relationship between participants’ behavior and the BOLD signal in RPE-and PRPE-sensitive regions, we also performed a simple regression analysis using the proportion of free-choices in which participants selected the PS+ key as a covariate. All results are reported at a threshold of p<.05 family-wise error (FWE) corrected within a small ventral striatal volume defined *a priori* based on an anatomical mask of the bilateral Nucleus Accumbens derived from the probabilistic atlas of Hammers et al. (Hammers et al., 2003).

## Results

### Behavior

Participants presented on average a preference for the PS+ key (M = .58, STD = .14), which was significantly above the chance level of 0.50 (*t*(19) = 2.46, *p* = .02) (Fig. 3A). This result is particularly interesting given that both keys were strictly equivalent in terms of final reward probability. However, there was a large variability across participants, with some participants presenting a clear bias towards the PS+ key and others presenting no clear preference for either key. This preference for the PS+ key was acquired throughout the task as suggested by a clear linear effect of time (*F*(1,19) = 8.66, *p* < .01, Fig. 3B).

**Figure 3:**
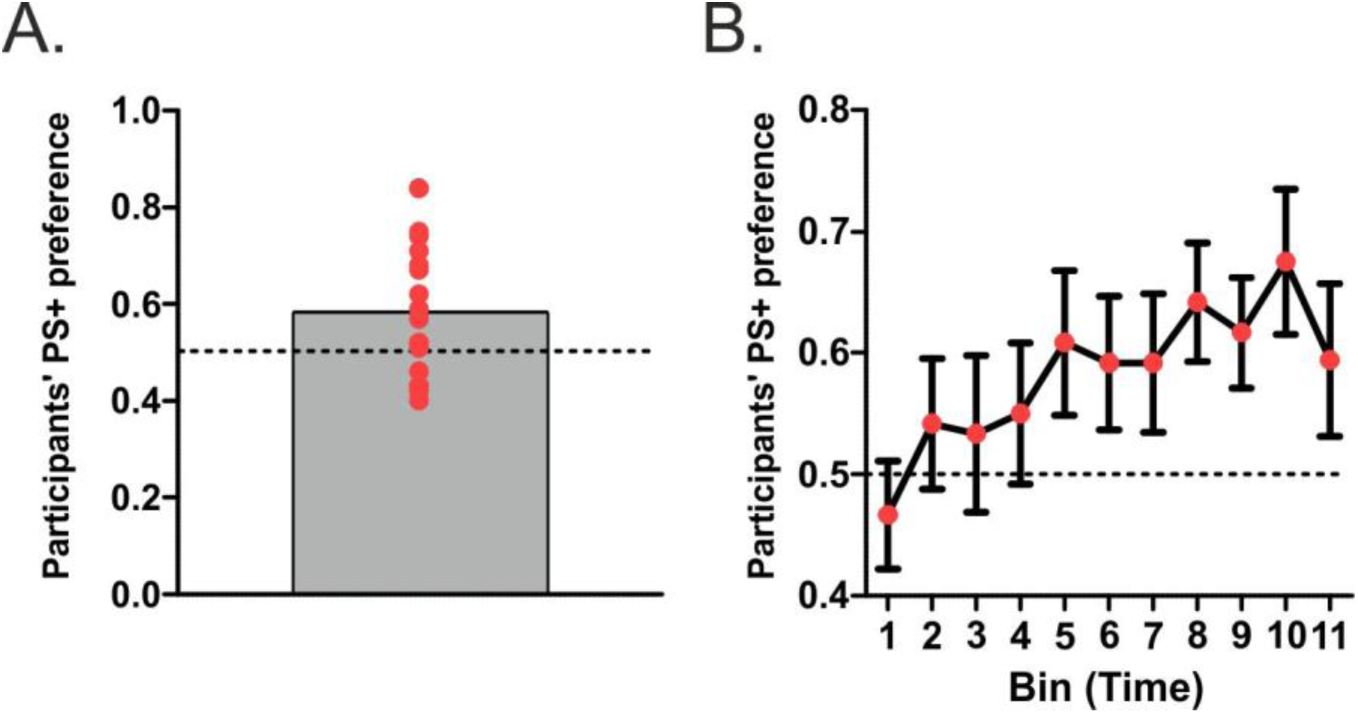
Behavioral results. **A)** Proportion of choices of the most pseudo-rewarding key (PS+). The grey bar represents the group average and red dots represent individual data points. **B)** Participants’ preference across trials (mean ± SEM). Each bin comprises 6 trials. Note a clear increase from the initial trials to the last trials of the task, demonstrating that the preference for the PS+ is acquired through learning.

The logistic regression analysis revealed a significant effect of both the history of rewards (*t*(19) = 2.3, *p* = .03) and pseudo-rewards, (*t*(19) = 2.6, *p* = .02) on participants’ choices, suggesting that participants were influenced by both types of feedback. As expected, the beta parameter quantifying the weight of pseudo-rewards in the choice process showed a strong correlation with participants’ preference: the strongest the preference for the PS+ key, the strongest the influence of pseudo-rewards in participants’ choices (rho(20) = .94, *p* < .001).

### Striatal representation of prediction errors

Figure 4 shows that brain activity in the bilateral ventral striatum correlated significantly with RPEs at the time of both the pseudo-feedback (RPE1, *p* < .001 FWE SVC; left: x=-10, y=9, z=-10; right: x=9, y=9, z=-7) and the final feedback (RPE2, *p* < .001 FWE SVC; left: x=-10, y=9, z=-10; right: x=9, y=9, z=-7). Ventral striatal activity also correlated with PRPE at the time of the pseudo-feedback (*p* < .01, FWE SVC; left: x=-9, y=9, z=-7; right: x=9, y=9, z=-7). The conjunction of the three contrasts (PRPE, RPE1 and RPE2) revealed significant activation in both the left and right ventral striatum (*p* < .01, FWE SVC; left: x=-7, y=9, z=-7; right: x=9, y=9, z=-7).

**Figure 4:**
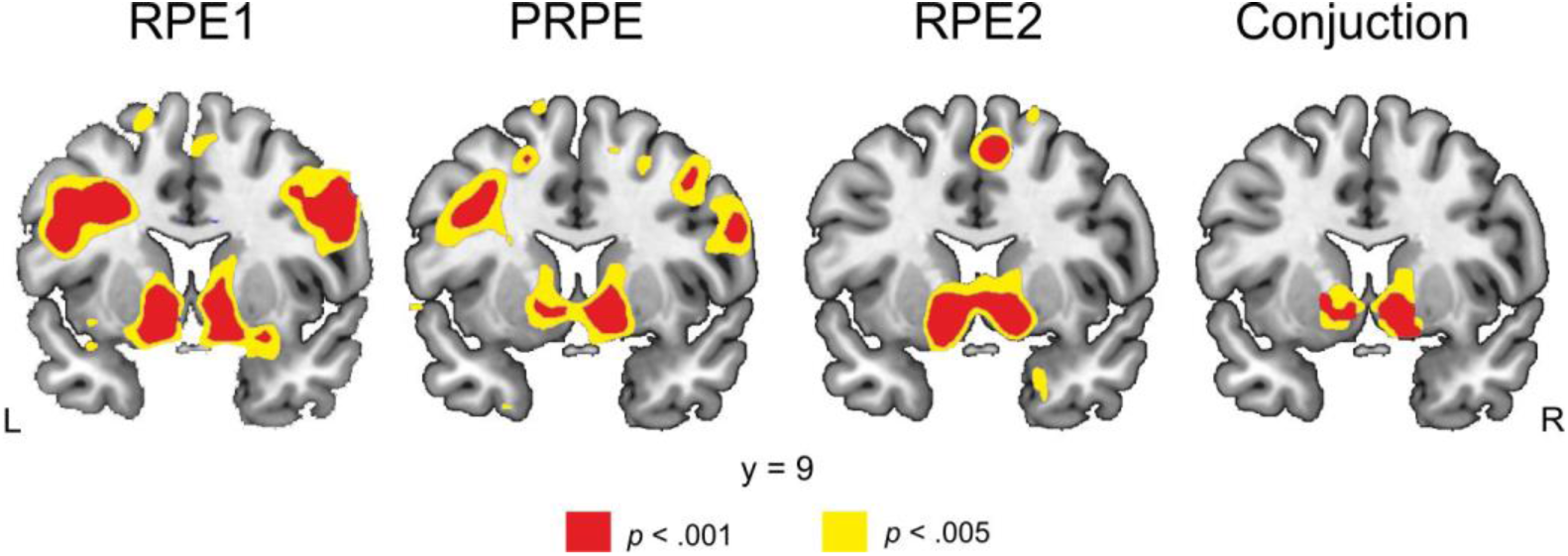
Striatal representation of reward and pseudo-reward prediction errors. T-maps showing the brain regions in which a positive relationship was observed across trials between BOLD activity and the various errors terms, namely RPE1 (reward prediction error at the time of pseudo-feedback), PRPE (pseudo-reward prediction error at the time of pseudo-feedback) and RPE2 (reward prediction error at the time of final feedback). The last panel on the right illustrates the conjunction of the three previous relationships in the ventral striatum. Note that, in all T-maps, peak activations in the bilateral ventral striatum survive a threshold of p<.01 family-wise error (FWE) corrected within a small volume defined *a priori* based on an anatomical mask of the bilateral Nucleus Accumbens (Hammers et al., 2003).

Since the ventral striatum is encoding both PRPEs and RPEs, we wondered whether the relative sensitivity to these two error terms would predict the behavioral preference for the PS+ key across participants. To assess this relative striatal sensitivity to PRPEs vs RPEs, we contrasted the corresponding parametric regressors reflecting how steeply these error terms scale with BOLD signal, and used the preference for the PS+ key as a covariate in a whole-brain simple regression analysis. We observed that the relative sensitivity to PRPEs vs RPEs was positively correlated with participants’ preference specifically in the left (*p* < .001 FWE SVC; x=-10, y=9, z=-7) and the right ventral striatum (*p* < .05 FWE SVC; x=9, y=6, z=-7). In other words, the higher the preference for the PS+ key, the greater the sensitivity to PRPEs compared with RPEs in the ventral striatum (Figure 5).

**Figure 5:**
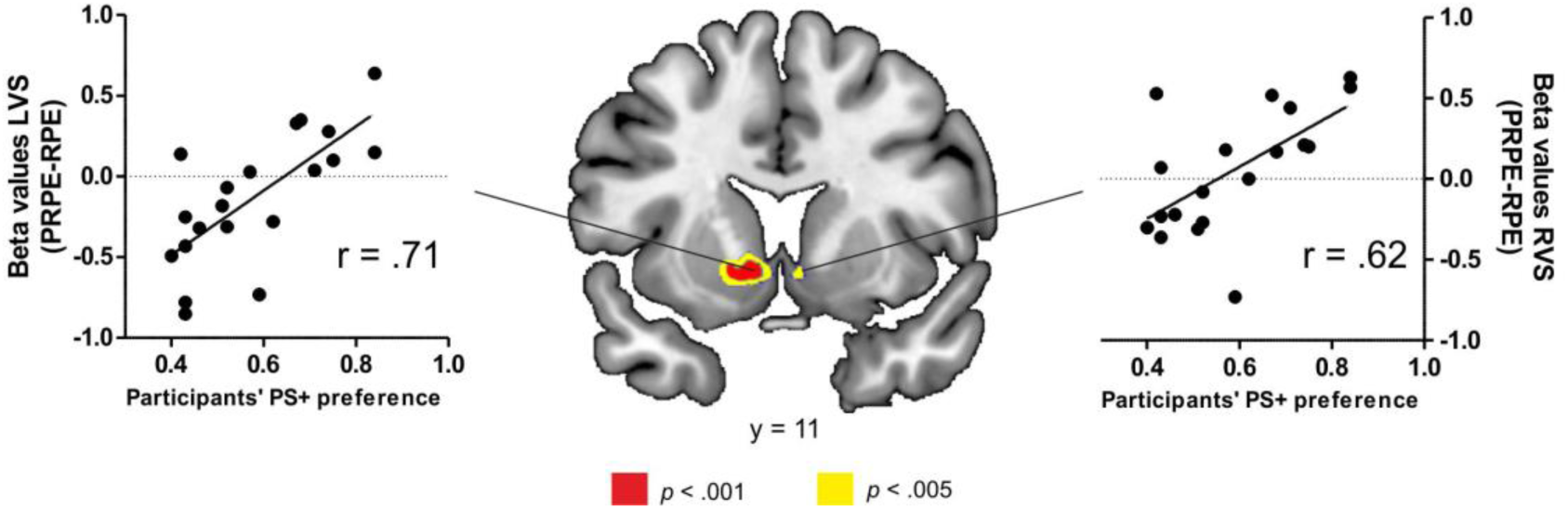
Brain-behavior relationship. T-map displaying the striatal voxels in which a significant relationship was found between relative striatal sensitivity to PRPEs vs RPEs and the behavioral preference for the PS+ key across participants. Note that peak activations in the bilateral ventral striatum survive a threshold of p<.05 family-wise error (FWE) corrected within a small volume defined *a priori* based on an anatomical mask of the bilateral Nucleus Accumbens (Hammers et al., 2003). The scatter plot, shown for purely illustrative purposes, illustrates the same relationship in the striatal voxels found to be significant in the voxel-wise analysis.

### Control experiment

We performed an additional behavioral experiment to rule out the possibility that the observed behavioral bias was merely driven by participants’ prior experience with keys and locks. Given that the sequence key-lock-open is quite common in everyday life, opening boxes may already have a pre-learned reward value that could explain the observed preference for the pseudo-rewarding key. To rule out this explanation, we tested a new group of 19 participants engaging in the same task as the fMRI experiment, but now using arbitrary cues instead of keys and locks (Figure S1, more details in Supplementary Information). Notably, participants also developed a preference towards the most pseudo-rewarding option (*t*(18) = 2.6, *p* = .018, Figure S2). These results replicate our previous findings using a more arbitrary design in a new group of participants, and thus confirm that participants’ bias towards pseudo-rewards is not driven by pre-learned reward values.

## Discussion

In the present study, we aimed to investigate whether pseudo-rewards could bias choice behavior-in a context where obtaining these pseudo-rewards does not confer any advantage– and whether such a bias might reflect striatal sensitivity to pseudo-rewards. In order to test these hypotheses, we developed a novel fMRI task in which participants had to first accomplish a sub-goal (unlocking a box = pseudo-reward) to obtain a probabilistic monetary reward. Participants presented a significant preference towards the key that was unlocking more boxes (i.e. more pseudo-rewarding) even though it did not lead to more monetary reward. Importantly, the inclusion of forced-choice trials ensured that participants sampled both options equally, and thus that the behavioral preference did not result from imbalanced learning between the two arms of the task (Niv et al., 2002). Remarkably, we replicated the same effect in a second group of participants engaging in a modified version of the task using arbitrary cues instead of keys and locks; this suggests that the observed bias towards the most pseudo-rewarding option is not driven by pre-learnt reward values. At the brain level, we observed a parallel representation of reward and pseudo-reward prediction errors in the ventral striatum. Notably, the relative striatal sensitivity to PRPEs vs RPEs predicted individual differences in the behavioral preference for the most pseudo-rewarding key. Our findings reveal a clear bias in participants’ choices driven mainly by the availability of pseudo-rewards. However, this bias cannot be considered sub-optimal in the present study since both options were equally rewarded and the bias was thus not associated with a cost. Nevertheless, it provides evidence that individuals tend to rely on pseudo-reward information in order to guide their choices during learning.

Our fMRI results showed that both RPEs and PRPEs are simultaneously encoded within the ventral striatum. This finding is consistent with the idea that the striatum is critical for computing prediction errors at several levels of complexity, and that dissociable prediction errors are integrated in this same structure (Daw et al., 2011, Diuk et al., 2013). In particular, our findings are in line with those previously reported by Diuk and colleagues (2013). This is remarkable given the differences in design between studies and the stringent orthogonalization procedure we used, which ensured that the PRPE regressor explained a significant amount of unique BOLD variance over and beyond the classic RPE. Additionally, Diuk et al. (2013) used a RL model in which RPEs only arose at the end of a trial when the outcome was presented. According to their model, reward expectancies did not vary from the first decision until the final outcome. In the context of the current task, their model would not anticipate potential outcomes based on whether the box was unlocked or not. In contrast, we have assumed a TD perspective, that is, a continuous-time model of learning in which any information between the first decision and the final outcome can lead to a RPE and can be used to generate new reward expectations.

Remarkably, the relative striatal sensitivity to PRPEs vs RPEs predicted participants’ behavior. Specifically, greater relative striatal sensitivity to PRPEs vs RPEs was associated with stronger preference for the most pseudo-rewarding key. In an analogous fashion, Daw et al. (2011) have shown that striatal sensitivity to model-free vs model-based RPEs signals predicted participants’ preference for model-free vs model-based learning strategy. Together, these results not only suggest that the ventral striatum encodes simultaneous learning signals, but also that these learning signals play a strong role in shaping behavior. This line of reasoning agrees with previous studies suggesting that the ventral striatum does not have an exclusive role in encoding reward prediction errors, but is involved in encoding the behavioral relevance of events (Delgado et al., 2008). For instance, Klein-Flugge et al. (2011) have shown that, in contrast to midbrain activity reflecting RPE signals in the absence of a behavioral policy, the ventral striatum responds mainly to those events that are relevant to guide behavior. Some authors have further theorized that striatal activity might mediate the dynamic attribution of incentive salience to events, causing them and their associated actions to become more relevant (Berridge, 2007).

It might be noted that the present results could also reflect sign-tracking behavior. Sign-tracking refers to the propensity of individuals to direct their behavior towards a stimulus that has become attractive and desirable as a result of a learned association between that stimulus and a reward (conditioned stimulus, CS), even when the location of the CS is not co-localized with the eventual reward (Flagel et al., 2009). Thus, the stimulus does not only act as a predictor of reward but also acquires motivational properties, becoming a “motivational magnet”. Previous studies have shown that there are large individual differences in this type of behavior, with some individuals engaging almost exclusively with the CS while others use this information to predict the location of the reward (goal-tracking) (Flagel et al., 2009; Robinson et al., 2014; Morrison et al., 2015). Similarly, in the current task, some participants may have assigned motivational properties to the pseudo-reward while others may have not. Thus, pseudo-rewards may also induce sign-tracking behavior and may, in part, explain the large individual differences in solving complex structured tasks in our everyday life. However, this account remains speculative, and further studies comparing performance in well-established sign-tracking paradigms with participants’ preference in the current task are required to better understand the relationship between pseudo-reward and sign-tracking behaviors.

It is important to emphasize that the behavioral bias towards the PS+ key cannot be explained by the desire to reduce uncertainty as reported in theories such as “temporal resolution of uncertainty” (Kreps and Porteus, 1978) or “information seeking” (Bromberg-Martin and Hikosaka, 2009). Previous studies have shown that animals avoid be uncertain about future outcomes (Bromberg-Martin and Hikosaka, 2009) and often prefer to have this uncertainty resolved earlier rather than later (Eliaz and Schotter, 2007). However, in the present study, the option leading to the biggest reduction in final outcome uncertainty was the least pseudo-rewarding key, which led to a fully predictable outcome (no-reward following locked box) in 70% of the cases. Thus, choosing the PS-key provided more advance information about the upcoming outcome than the PS+ key, which was associated with full uncertainty resolution at the time of pseudo-feedback in only 30% of the trials.

In sum, we have shown that pseudo-rewards can influence participants’ choices independently of the reward value associated to them. Thus, although they are critical for speeding up learning in complex environments, they might potentially lead to irrational behavioral biases. Additionally, using a novel paradigm we have shown that RPEs and PRPEs are simultaneously encoded in the ventral striatum, and that the relative sensitivity of the striatum to PRPEs vs RPEs predicted participants’ choices, emphasizing the relevance of the striatum in complex learning problems and decision-making.

